# Characterization of the hemodynamic response function in default mode network subregions for task-evoked negative BOLD responses

**DOI:** 10.1101/2025.09.19.677202

**Authors:** Hengda He, Bardiya G. Yazdi, Amirreza Sedaghat, Seyed Hani Hojjati, Sindy Ozoria, Peter Chernek, Jenseric Calimag, Farnia Feiz, Qolamreza R. Razlighi

## Abstract

While the hemodynamic response function (HRF) of the positive blood oxygenated level dependent (BOLD) signal is well characterized in the functional magnetic resonance imaging (fMRI) field, studies investigating the shape and properties of the negative BOLD HRF, often observed in the default mode network (DMN) regions, are rare. In this study, we first investigated the linearity of the task-evoked negative BOLD response (NBR) with respect to stimulus duration. Then, we obtained the shape of the HRF from each region of the DMN, using an in-house developed unbiased and robust iterative deconvolution technique. Our results demonstrated that each region of the DMN represented a unique HRF, which was not only substantially different from the HRF of the positive BOLD signal, but also different from the HRF extracted from the other DMN regions. We replicated these findings using different fMRI datasets with distinct task paradigms. When comparing the HRF across DMN sub-regions and across tasks, our results demonstrated a significantly higher inter-regional variability compared to inter-task variability. Furthermore, our performance correlation analysis illustrated that the HRFs in the DMN sub-regions extracted by our iterative FIR method were not biased by the task performance level. Altogether, we demonstrated the linearity of the NBR, introduced a robust HRF extraction approach to obtain a distinct HRF for each DMN sub-regions, which were different from the HRF of the positive BOLD signal. The results suggest there is possibility that each node of the DMN might have different underlying neural and/or vascular mechanisms.

## Introduction

Brain’s neuronal activity is often compensated by a disproportionally large increase of the local cerebral blood flow (excessive hyperemia) through a cascade of events referred to as *neurovascular coupling* (Gordon, Mulligan, and MacVicar 2007; Iadecola and Nedergaard 2007). This large increase in local cerebral blood flow, and the subsequent rise in blood volume, alters the T_2_ relaxation time in the activated regions. These changes, in turn, cause alterations in the functional magnetic resonance imaging (fMRI) signal received from that region (Buxton et al. 2004). This change in the MR signal is commonly defined as blood oxygenation level dependent (BOLD) signal (Huettel, Song, and McCarthy 2014; Ogawa et al. 1990). The shape of this signal change due to a brief spike of neural activity is called the hemodynamic response function (HRF) of the BOLD signal. The shape and other properties of the BOLD response have been investigated in numerous studies (Boynton et al. 1996a; Clark, Maisog, and Haxby 1998; Courtney et al. 1997; Dale and Buckner 1997; Friston et al. 1994) which demonstrated a sluggish and delayed increase in MR signal in comparison to baseline period that reaches to its peak magnitude at 6 to 7 seconds after the termination of the external stimuli. After its peak, the MR signal starts decaying back to its baseline in almost the same amount of time followed with an extended period of undershoot (between 10 to 15 second), adding up to about 25 seconds of continuous changes in MR signal even for a brief stimulus as short as 500 milliseconds. In contrast, the MR signals from other brain regions, often unrelated to the stimuli, are shown to have a change in opposite direction where a decrease in MR signal with similarly delayed and slow-paced change in about the same period of time (8 to 14 seconds) is observed when compared with baseline period. This reduction in MR signal is also often followed by an extended overshoot (Lustig et al. 2003). We, henceforth, refer to this inverse BOLD signal as negative BOLD response (NBR), and similarly call the increase in MR signal as positive BOLD response (PBR) in this work.

By late 1990s, using positron emission tomography (PET), Shulman and colleagues reported that during a broad range of goal-oriented tasks a constellation of brain regions appears to lose blood flow when compared to passive conscious rest (Shulman et al. 1997). Such findings, few years later, led to the introduction of “default mode” of the brain by Raichle which asserts that there are functionally interconnected brain regions that are active during processes that usually happen during rest, and their activities are temporarily suppressed during engagement in any external and attention demanding tasks (Raichle et al. 2001). These regions include large segments of lateral parietal cortex (i.e. angular gyrus and tempro-parietal junction), the medial parietal cortex (i.e. posterior cingulate and precuneus), ventro-medial and superior frontal cortex, middle temporal lobe and hippocampus. Despite an overwhelming number of studies reporting NBR from DMN regions during variety of tasks, little is known about their magnitudes, dynamics, and other properties of their hemodynamic response function (HRF). For instance, while there are studies reporting that the task load modulates the magnitude of the NBR in the DMN (Greicius et al. 2004; Mayer et al. 2010; McKiernan et al. 2003; Todd, Fougnie, and Marois 2005; Tomasi et al. 2009), the NBR linearity with respect to stimulus duration, to our knowledge, has not been investigated. This essentially raises a major concern about all findings with NBR using general linear modelling analysis, which is the most common technique used for analyzing BOLD fMRI. Therefore, in this study we first start by examining the linearity of the NBR with respect to stimulus duration in the main nodes of the DMN.

There is also no evidence that the HRF of the NBR has the same shape and dynamic as the commonly used HRF (double-Gamma) for PBR. In fact, few studies that have investigated the time-course of the NBR from the DMN regions (in comparison with PBR) have shown significant differences between them (Farooqui and Manly 2018; Lustig et al. 2003; Mayer et al. 2010; Meltzer, Negishi, and Constable 2008). Furthermore, while the initial hypothesis of the DMN was not concern about the regional heterogeneity of the NBR observed from different DMN regions, more recent studies (Leech et al. 2011; Mayer et al. 2010; Shulman et al. 2007) highlight the possibility that each region of the DMN may have distinct roles in the brain functionality thus representing a distinct temporal and/or spatial characteristics in its BOLD responses. Therefore, we next extract the shape of the HRF from each region of the DMN separately. We hypothesise that the HRF from different regions of the DMN will demonstrate distinct dynamic/shape highlighting the possibility of having separate underlying neuronal and/or vascular mechanisms as well as having different roles in the brain functional orchestration.

The role of the DMN as a whole and specific functionality of its nodes in the brain is controversial and still under investigation. However, alterations of the NBR in the DMN regions have been shown in normal aging and numerous clinical populations including but not limited to Alzheimer’s disease (Ghaderi Yazdi and Razlighi 2025; Lustig et al. 2003; Persson et al. 2007), mild cognitive impairment (Threlkeld et al. 2011), autism (Spencer et al. 2012; Vogan et al. 2018), schizophrenia (Hanlon et al. 2016; Pomarol-Clotet et al. 2008), attention deficit and hyper activity disorder (Fassbender et al. 2009; Liddle et al. 2011; Metin et al. 2015), and among others. All these studies assumed, without any scientific evidence, that the spatial and temporal characteristics of the NBR signal is similar to that of the PBR. While this unvalidated assumption could potentially generate acceptable results in some studies and applications, it can also generate false-positive findings that could easily be mistaken as a genuine breakthrough. For example, regional differences in how much each subregion of the DMN deactivates could be explained by variations in the underlying properties of each region, such as the regional specificity of the HRF or the linearity in the regional responses to task stimuli durations. Furthermore, brain-behavior relationship is often reported using NBR from the DMN; However, using erroneous HRF could not only reduce the sensitivity of the analysis to detect true underlying effects, but also could generate false regional differences in the brain-behavior relationships. Therefore, it is of paramount importance that the spatial and temporal characteristics of the NBR be investigated and fully understood to be able to make any meaningful interpretation about the results obtained from NBR in the DMN regions.

Existing reports by our group and others indicate that the magnitude of task-evoked NBR are smaller than PBR (He et al. 2022; Lustig et al. 2003; Razlighi 2018; Shulman et al. 2003). Thus, using a conventional method developed for investigating PBR may not generate an accurate and reproducible result for the NBR extracted from the DMN regions. This might be the reason that studies investigating the spatial and temporal characteristics of the NBR are so rare in the field. To address this issue, we have developed and implemented an unbiased and iterative finite impulse response (iFIR) method to extract an accurate and reproducible HRF for the NBR detected from the DMN regions (Goutte, Nielsen, and Hansen 2000). Extraction of an unbiased kernel that represents the underlying hemodynamic response in each anatomical region of the DMN not only improved the sensitivity of the analysis to detect the actual brain-behavior relationship but also helps direct the investigations targeting the underlying neural and/or vascular substrates of the NBR at each node of the DMN.

In this paper, we start by investigating the linearity of the NBR with respect to stimulus duration in the main nodes of the DMN. We then use our developed iFIR deconvolution method to extract the HRF kernel from sub-regions of the DMN and investigate their timing and magnitude differences aiming to demonstrate heterogeneity in the shape and dynamic of each DMN region’s HRF. To establish robustness of our iFIR method and the obtained HRF kernels, we replicate our findings using fMRI data with various task paradigms. Lastly, we investigate the brain-behavior relationship by relating the task performance (accuracy and reaction time) to the amplitude of the HRF in each DMN sub-region. Our results demonstrated that the HRFs in the DMN sub-regions extracted by our iFIR method are not biased by the task performance. Our findings indicate that each region of the DMN presents distinct HRF kernel which highlight the possibility of having unique underlying neural and/or vascular mechanism specific to that region.

## Material and Methods

### Participants

Our dataset included 61 young, healthy, right-handed participants (age = 25 ± 3.5 years, male/female = 35/20) recruited from 10-mile radius of the Weill Cornell Medicine campus using random market mailing. All participants in the study signed an informed consent form before the scanning session and were compensated for their time spent in the study. The research experiments were performed in accordance with relevant guidelines and regulations approved by the Weill Cornell Medicine Institutional Review Board.

### fMRI experimental design

We used 6 different cognitive tasks tapping into four different cognitive domains (perceptual speed, crystallized memory, fluid reasoning, and episodic memory). All tasks were designed using ePrime 3 (https://pstnet.com/products/e-prime/) and employed an event-related fMRI paradigm to be able to optimally investigate magnitudes and dynamics of the HRF extracted for each of the DMN regions. All fMRI experiments were designed visually and presented within a square/rectangular aligned to the horizontal meridian and were projected to a translucent screen located at the far end of the scanner. Participant were lying supine inside the magnet bore, wearing noise isolating MR safe earbuds, and were seeing the screen using a mirror located on the head-coil. Subjects were first trained outside of the scanner to learn and perform the task comfortably and accurately in a short training session outside the scanner (∼30 minutes). All subjects learned the task correctly. All responses were made using a button press device (https://pstnet.com/products/celeritas/) with five buttons. Across all tasks administered during the training session as well as the tasks administered during the actual fMRI scanning session, the thumb corresponded to choice number one, index finger with choice two, middle finger with choice three, ring finger with choice four, and little finger with choice five. Once the subjects made their choices, the stimulus was removed from the screen, and a background white screen was presented until the next stimulus appeared. All scan durations were set to 10 minutes to control for the variability due to the scan duration. Due to the differences in the difficulty of each cognitive task, the number of trials could not be balanced across all tasks. Instead, the duration in which participants are engaged in the task performance versus the rest period was balanced for all tasks (using the median response time, the ratio of task/rest periods was set to 0.5 for each task).

Next, we explain each task design paradigm briefly.

#### Synonym

This is a crystalized memory task where a probe word is presented to the participant at the top of the screen in all capital letters and four choice words were listed underneath. Figure 1 shows the timing used for this task. Subjects were to select the word most similar in meaning to the probe word. In total, 40 trials were presented to each participant, and participants were given 10 seconds to make a choice. The onsets of trials are jittered using a continuous uniform distribution in the range of 0-20 seconds. 54 out of the 61 participants finished synonym task-based fMRI scan.

**Figure 1.**
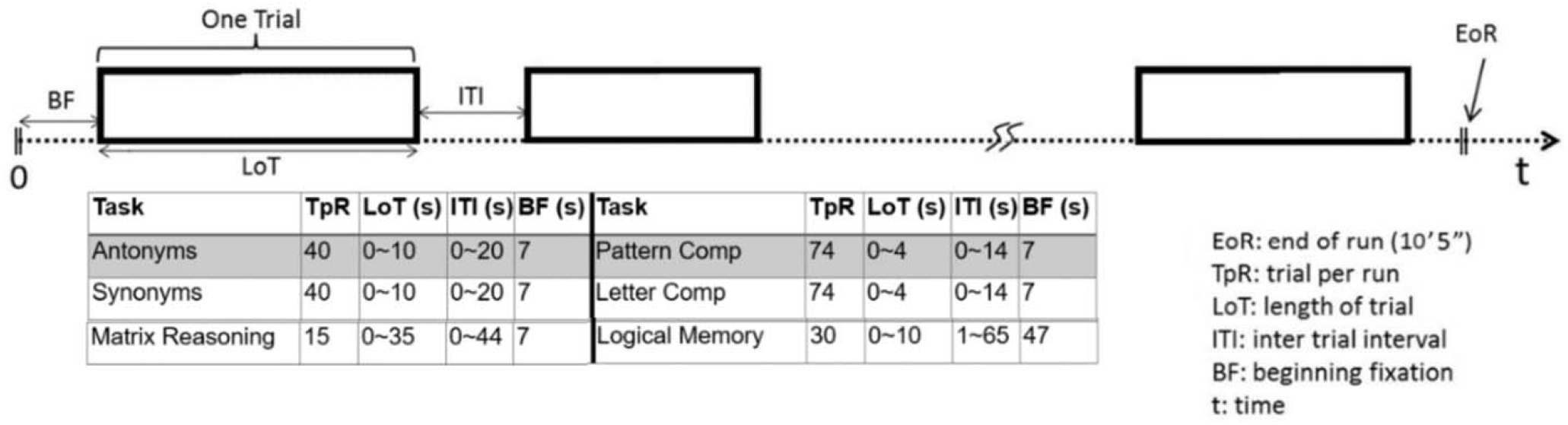
Illustration of a segment of the time-course in the event-related tasks. Each trial started with a beginning fixation period (BF), followed by the task presentation with a length of trial duration (LoT). Each subject was scheduled to participate in six tasks in the MRI scanner, where parameters of each task design are illustrated in the table.

#### Antonym

This is another crystalized memory task very similar to synonym task, and as it appears from the name, the only difference is that, in this task, participants were instructed to match a probe word to the word most different in meaning. All the timings and other design aspect of the task are the same. 55 out of the 61 participants finished antonym task-based fMRI scan.

#### Pattern comparison

This is a perceptual speed task where two abstract figures, consisting of a varying number of lines connecting at different angles, are presented alongside one another. Participants are instructed to indicate whether the presented figures are identical by using a differential button press. Specifically, participants were instructed to press the button aligned with their right thumb if the figures were identical or their right index finger if the figures were not identical. The maximum allowable time to make a response was four seconds, and the onset of the stimuli was jittered by applying a randomized inter-stimulus interval (ISI) generated from a uniform continuous distribution in the range of 1.0 – 4.0 seconds. The events were randomized once, and the same random sequence was used for all the subjects. Participants were administered one run consisting of 74 trials completed in 10 minutes. The timing of this task throughout the whole 10-minutes scan time are shown in Figure 1. Participants were instructed to respond to each stimulus accurately and as soon as possible. 57 out of the 61 participants finished pattern comparison task-based fMRI scan.

#### Letter comparison

This task is another perceptual speed task where instead of two figures, two strings of letters of equal length were presented to the participants. Participants were instructed to identify whether the two sets of letters were identical. Similar to pattern comparison, participants needed to press the button aligned with their right thumb if the letters were identical or their right index finger if the figures were not identical. All other aspects of this task were the same as pattern comparison task explained above. 54 out of the 61 participants finished letter comparison task-based fMRI scan.

#### Matrix reasoning

This is a fluid reasoning task which is a simplified version of the famous Raven Matrix Reasoning (Raven, Raven, and Court 1962). In this task eight patterned figures are presented in the eight cells of a 3×3 matrix leaving the bottom-right cell empty. There are rules applied to rows and columns. Participants were to identify the rules and find out what should be the patterned figure in the missing cell. Five choices are given underneath the matrix, and participants were to choose the most appropriate choice that completed the matrix. In total, 15 trials were presented during a 10-minute scan and participants are given ample time to make their decision (35 seconds). The onset of the trials was jittered by adding a random ISI time selected from a continuous uniform distribution in the range of 0 - 44 seconds. 56 out of the 61 participants finished matrix reasoning task-based fMRI scan.

#### Logical memory

This is an episodic memory task which measures the number of details recognized within stories. Subjects are required to remember all details from stories presented on the computer screen, then asked to answer detailed multiple-choice questions about the story, with four possible answer choices. The stories were presented one at a time, and participants had 30 seconds to read each story. Next, participants retained the information for 10 seconds before the first probe question appears on the screen. For each story, 10 probe questions were presented resulting in 30 probe questions in total. The onset of the encoding stories and probe questions were jittered using a continuous uniform distribution in the range of 1 to 65 seconds. Four answers were listed beneath each probe question and participants were instructed to choose the most appropriate answer by pressing a button associated with the number of the choice presented on the screen. 52 out of the 61 participants finished logical memory task-based fMRI scan.

### MRI acquisition parameters

All MRI scans were carried out in two sessions separated with at least one week apart. The order of the tasks in the two sessions were arranged in a way that the two fMRI tasks scans belonging to the same cognitive domain were always performed in different sessions. A research dedicated 3 Tesla Siemens Magnetom Prisma scanner equipped with a 64-channel head-coil, and 80 mT/m gradient system was used in this study with a multiband T_2_*-weighted echo-planar imaging (EPI) pulse sequence [TR/TE = 1008/37 ms; flip angle = 52^°^; FOV= 208 × 208 mm; matrix-size = 104 × 104; voxel-size = 2 × 2 × 2 mm; 72 axial slices; multiband factor = 6]. The duration of fMRI scan was 10 minutes and 5 seconds (600 volumes). The phase encoding direction (posterior - anterior) was alternated for fMRI scans, to be used retrospectively for *TopUp* geometric distortion correction. An accompanying T_1_-weighted magnetization-prepared rapid gradient-echo (MPRAGE) structural image was collected [TR/TE = 2400/3 ms; flip angle = 9^°^; FOV= 256 × 256 mm; matrix-size = 512 × 512; voxel-size = 0.5 × 0.5 × 0.5 mm; 320 axial slices] for localization and spatial normalization of the functional data in each participant.

### Structural and functional MRI preprocessing

All participant’s structural T1-weighted MRI scans were processed using FreeSurfer software package (https://surfer.nmr.mgh.harvard.edu/) to get the parcellated cortical and segmented subcortical regions according to Desikan Killiany atlas (Desikan et al. 2006). Functional images were preprocessed using an in-house developed pipeline, with the aim to optimize data quality and address challenges associated with multiband EPI sequences. All analyses in this project were performed in the subjects’ native space to prevent inaccuracies due to spatial normalization (Liu et al. 2017; Razlighi et al. 2014; Seibert and Brewer 2011). The preprocessing pipeline began with spatial realignment using FSL’s *mcflirt* algorithm. A single-band reference (SBRef) scan was used as the reference volume for rigid-body registration to ensure precise alignment of functional volumes throughout the scan. Following realignment, slice timing correction was performed using FSL’s *slicetimer* tool to temporally align voxel timeseries to the beginning of the TR. Spatial smoothing was applied using FSL’s *susan* with a 5 mm full-width at half-maximum (FWHM) kernel.

To remove motion-related artifacts, ICA-based Automatic Removal of Motion Artifacts (AROMA) (Pruim et al. 2015) was employed. The motion parameters derived from the spatial realignment step were input into AROMA to help identifying motion-related independent components. AROMA also requires an accurate brain mask and a warping field that maps native space fMRI images to MNI template space. The fMRI native space brain mask was generated using the participant’s structural scan (MPRAGE) and corresponding FreeSurfer segmentation/parcellation. Specifically, the brain mask in the FreeSurfer structural space was transformed to the participant’s native fMRI space using the inverse of the rigid-body co-registration transformation and the inverse of the geometric distortion correction warping field obtained from FSL’s *topup* tool. For geometric distortion correction, an additional fMRI scan with the opposite phase encoding direction was used to estimate the warping field. For the registration between subject structural space to MNI space, non-linear registration was performed using ANTs. To generate a single composite warping field between subject’s native fMRI space and the MNI space, the non-linear warping field was then combined with the geometric distortion correction and the rigid-body co-registration transformation. The final warping field, along with the motion parameters and brain mask, was used by AROMA to identify and remove motion-related components from the data. Given evidence showing the robustness of task-based fMRI (tb-fMRI) to motion artifacts, a non-aggressive AROMA approach was employed.

After AROMA, global intensity normalization was applied to ensure a global median fMRI intensity of 10,000 (after outlier removal). Next, temporal filtering was performed, using a high-pass filter with a cutoff frequency f > 0.01 Hz. To eliminate any residual motion artifacts, scrubbing was performed. Specifically, scrubbing was implemented based on thresholds of framewise displacement larger than 0.5 mm, or global gray matter percent change signal had a RMSD larger than 0.75.

Five regions of interest (ROI) masks were generated based on FreeSurfer parcellations. We included three main nodes in the DMN: 1) Posterior cingulate cortex (PCC): posterior cingulate, isthmus cingulate, and precuneus regions; 2) Inferior parietal lobe (IPL); and 3) medial-orbito-frontal cortex (MFC). For comparison purposes, a binary mask of primary visual cortex (lateral occipital cortex; LOC) is also used to generate the results for PBR. We additionally included the middle temporal gyrus (MTG) and hippocampus (Hipp) as the sub-regions of the DMN to extract its regional HRF.

### Linearity of the NBR in respect to stimuli duration

To assess the linearity of the NBR with respect to stimulus duration in nodes of the DMN, we first analyzed fMRI data from the synonym task, which elicits robust task-related deactivation in the DMN (Ghaderi Yazdi and Razlighi 2025). Firstly, stimuli were grouped by duration: 2, 4, 6, 8, and 10 seconds. As illustrated in Figure 2, we modeled the task-evoked BOLD responses using three models: (1) no modulation, (2) saturated modulation, and (3) linear modulation. The “no modulation” model assumed that the fMRI response remained constant regardless of stimulus duration. To avoid bias associated with the assumption of specific HRF shapes, a Gaussian kernel (mean = 5 sec, standard deviation = 1.5 sec) was used instead of a double-gamma function. The “saturated modulation” model represented a simple non-linear scenario where stimulus with a longer duration evoked a response that saturated upon reaching a peak but maintained the same duration scaled to the stimulus length. The “linear modulation” model followed the most common assumption in the linear modelling analysis of fMRI data, where the evoked response was modeled as a linear convolution of the stimulus with the HRF kernel. In the “linear modulation” model, both the amplitude and duration of the evoked response scaled with the stimulus duration. Next, these models were fitted to the voxel-wise fMRI time series, and model fit was evaluated using R-squared values. We quantified the R-squared values in the most deactivated voxels in each DMN region (100 voxels). The differences in R-squared between models were calculated for each participant. We hypothesized that the “linear modulation” model would provide the best fit, with the highest R-squared values among the three models. To test this hypothesis, Student’s t-tests were conducted to determine if the differences in R-squared between the linear model and the other models were significantly greater than zero. The same approach was applied to evaluate PBR in the primary visual cortex.

**Figure 2.**
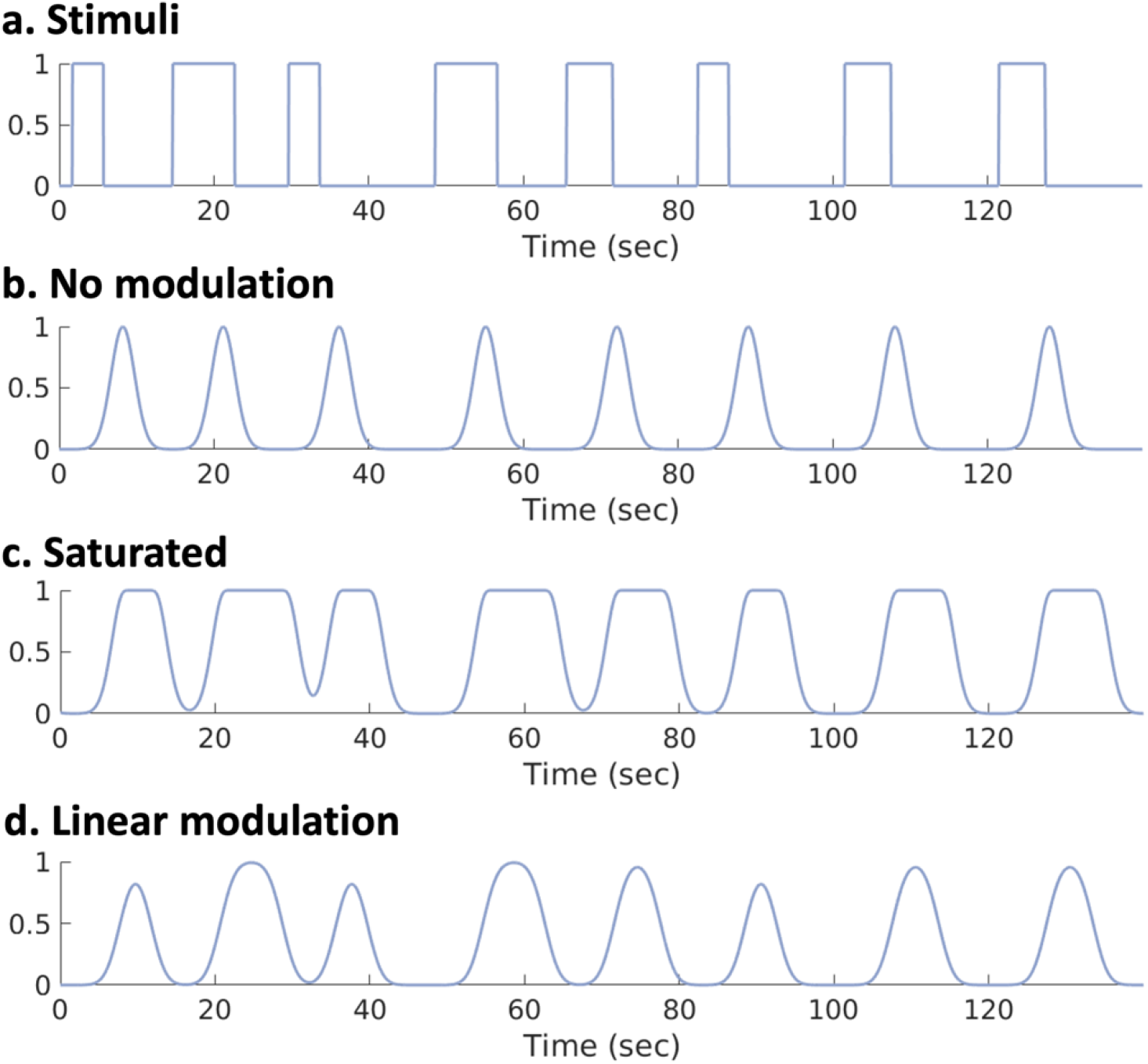
In the linearity analysis, task-evoked BOLD responses were modeled with three conditions including: no modulation, saturated modulation, and linear modulation. The figure illustrates a segment of the stimuli onset and offset (a), and the no modulation condition (b) assumed that the fMRI response remained constant regardless of stimulus duration. The saturated modulation condition (c) assumed a stimulus with a longer duration evoked a response that saturated upon reaching a peak but maintained the same duration scaled to the stimulus length. Lastly, in the linear modulation condition (d), the evoked response was modeled as a linear convolution of the stimulus with the HRF kernel, where both the amplitude and duration of the evoked response scaled with the stimulus duration.

### Unbiased iterative finite impulse response

This approach extracts a negative hemodynamic response from each anatomical region within the DMN in an unbiased manner. The algorithm is specifically designed to avoid any prior assumptions about the time course of responses from deactivated voxels in the brain. And this approach is optimized for obtaining subject-specific hemodynamic responses to improve model fitting. To achieve this, a FIR deconvolution method was applied to each voxel’s time-series within each sub-region of the DMN, using the stimulus paradigm to extract the HRF for the given voxel. Firstly, the initial kernel was set as the same Gaussian function that was used in the linearity analysis. The subsequent steps of the analysis were performed separately for each region. The initial HRF was used as a starting kernel in a GLM, generating activation maps for each participant. Then, voxels showing the strongest negative responses were selected (100 voxels), and their deconvolved HRFs were averaged across participants to create the kernel for the next iteration. This iterative procedure continued until the kernel converged, defined as when the squared difference between successive kernels was less than 10^−6^. Two additional DMN sub-regions of MTG and Hipp were included for this analysis. The same approach was applied to the fMRI signal in the primary visual cortex to extract HRF of PBR. Given the established understanding that the magnitude of NBR across the brain is generally weaker than PBR, instead of selecting the most activated/deactivated voxels, we used a threshold of t-score > 6 for extracting the HRF of PBR.

In a separate experiment, various features of HRFs from each DMN sub-region were extracted to compare responses across these regions. This analysis aimed to identify differences in the hemodynamic response characteristics between sub-regions of the DMN. The extracted features included onset time, time to peak, amplitude, time from peak to baseline, and time to overshoot. Onset time was defined as the point at which the HRF exceeded 10% of its peak amplitude. The time from peak to baseline was measured as the duration between the peak time and the point when the HRF returned to its estimated pre-stimulus baseline level. The time to overshoot was defined as the time required for the HRF to reach its maximum positive value after returning to baseline, starting from time zero. More details on HRF characteristics are available in a previous study (He et al. 2022).

### Behavioural Analysis

To examine the relationship between the HRF of NBRs and task performance of the participants, we used the subject-specific accuracy over median response time as a measure of performance. We employed Pearson correlation coefficients to assess the associations between task performance and the amplitude of the HRF in the sub-regions of the DMN. False discovery rate (FDR) was used for multiple comparison correction. The relationships were assessed again after controlling age and sex. We hypothesized that the extracted NBR HRFs should not be related to any difference in task performance.

## Results

### Linearity

The linearity of NBR was examined on the synonym task-based fMRI data. Specifically, fMRI time series in the main nodes of the DMN were fit with three models using GLM, where task-evoked BOLD response was modelled with different modulation assumptions: (1) no modulation; (2) saturated modulation; and (3) linear modulation. We expect the linear modulation model capture the most variance in the fMRI signal with the highest R-squared values. As shown in Figure 3, we reported the difference in R-squared values comparing the linear modulation modelling with the no modulation and the saturated modulation models. Linear modulation model achieved significantly higher R-squared values compared to the no modulation model in all three nodes of the DMN (IPL: p < 3.89×10^−11^; PCC: p < 1.28×10^−11^; MFC: p < 1.17×10^−09^), as well as LOC (p < 3.70×10^−11^). Similar results were obtained comparing linear and saturated modulation model (LOC: p < 6.41×10^−17^; IPL: p < 1.67×10^−13^; PCC: p < 2.00×10^−12^; MFC: p < 1.33×10^−08^). The results for visual cortex are also depicted for comparison purposes. These results provided evidence supporting the linearity assumption of task modulation in the main nodes of the DMN.

**Figure 3.**
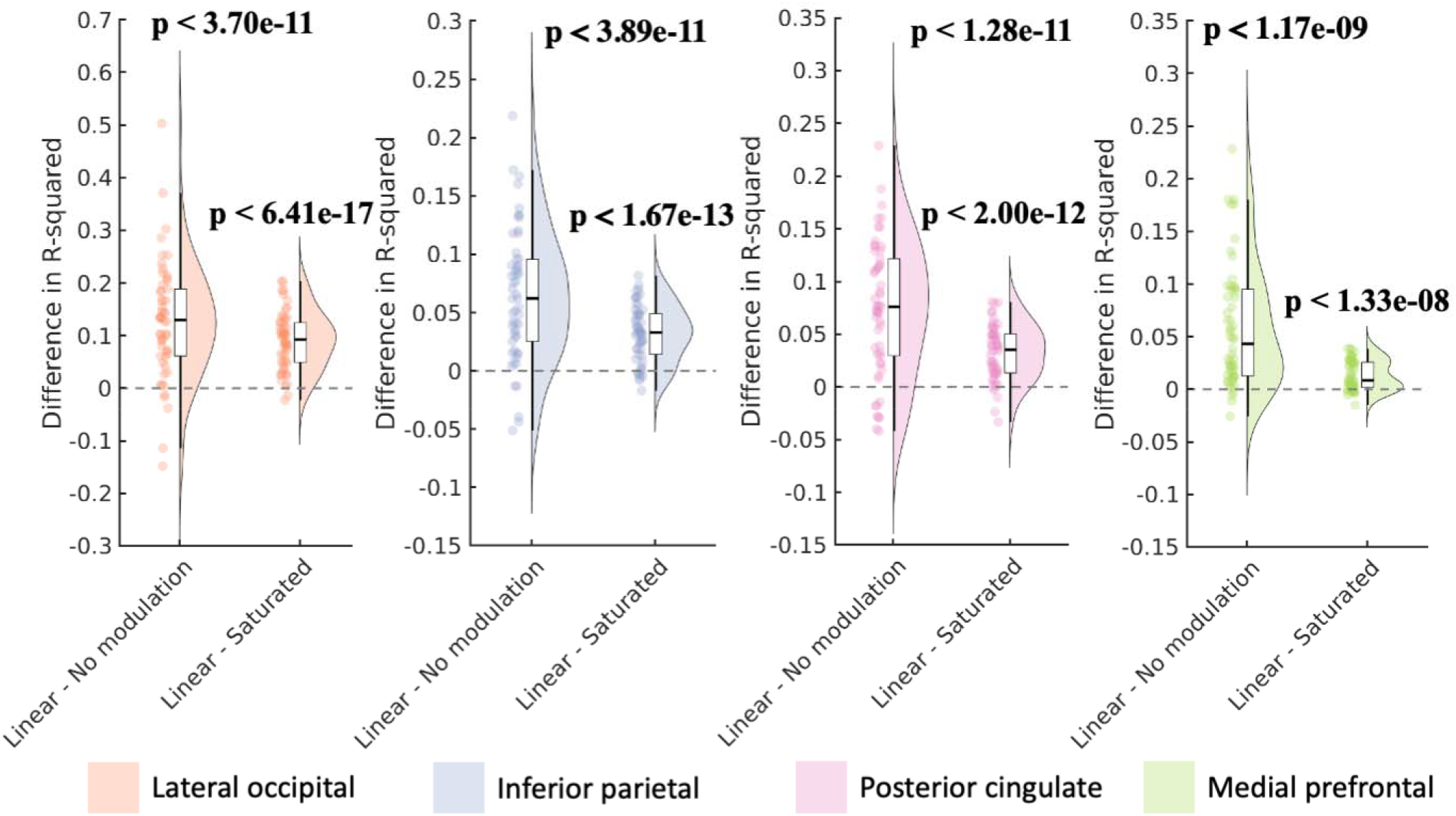
In the linearity analysis, we computed the difference in R-squared values comparing the linear modulation modelling with the no modulation and the saturated modulation models. In the synonym task, linear modulation model achieved significantly higher R-squared values compared to the no modulation model, as well as the saturated modulation model.

### HRF Extraction

To extract regional HRF, an unbiased iterative finite impulse response approach was employed for each sub-region of the DMN and the LOC. And two additional DMN sub-regions were included (Hipp and MTG). The shape of extracted HRF throughout the iterations and the initial HRF were illustrated for the synonym task data (Figure 4). Each region took a different number of iterations to converge (LOC: 3; IPL: 6; PCC: 6; MFC: 5; Hipp: 6; MTG: 5). To validate the HRF extraction method, we further applied the method to the other task-based fMRI data from the same and different cognitive domains including: antonyms, letter comparison, pattern comparison, matrix reasoning, and logical memory. When examining the task-evoked responses, different task timing design across these tasks might potentially confound the extracted HRF. Thus, we first assessed the relationship between the median task durations of the task and its corresponding HRF amplitude for the task. As shown in Figure 5a, our results demonstrated that the extracted HRF responses were not systematically influenced by the variation in task duration, indicating that our method is robust to differences in stimulus timing. In Figure 5b, we averaged the HRF of the NBR across different tasks for each sub-region of the DMN (error bar represents standard deviations across tasks). The strongest NBR was observed in the IPL, while the weakest NBR was observed in the Hipp. As shown in Figure 5d, we also illustrated the averaged HRF across DMN sub-regions (error bar represents standard deviations across DMN sub-regions) for each task. The strongest NBR was observed in the pattern comparison tasks, while the weakest NBR was observed in the logical memory task. Additionally, we performed Wilcoxon rank sum test between the variability in the HRFs across the sub-regions (inter-regional variability; see error bars in Figure 5d) and the inter-task variability (see error bars in Figure 5b), by pooling all the variance values between 1s to 10s after stimuli onset. Our results demonstrated a significantly higher inter-regional variability compared to inter-task variability (p < 8.36×10^−4^). Together, these results provide evidence that the HRF of the NBR from DMN does not change significantly during different cognitive task whereas different nodes of the DMN have slightly different HRF dynamics.

**Figure 4.**
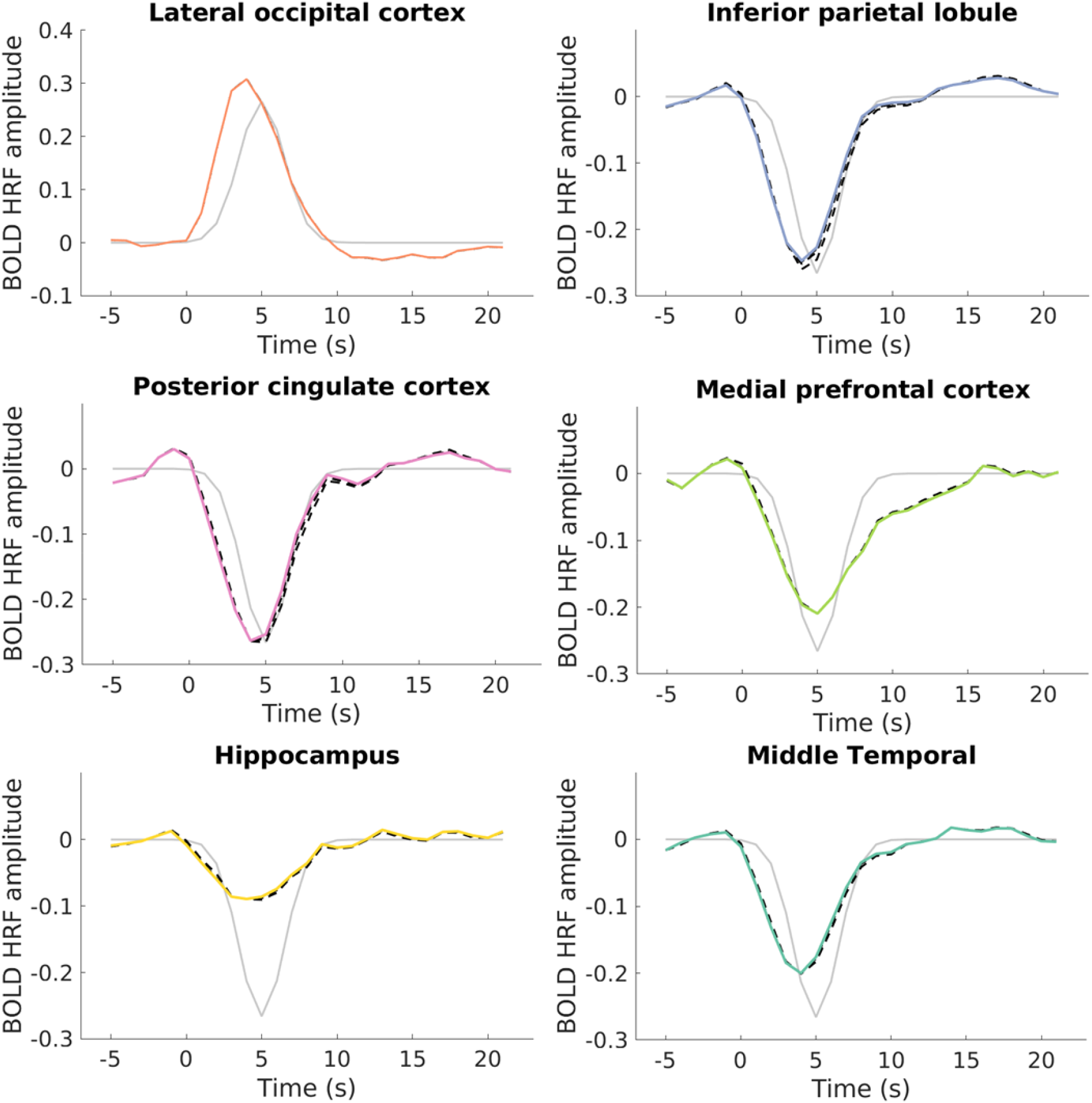
An iterative finite impulse response (FIR) method was used to extract hemodynamic response function (HRF) of the BOLD signal in each default mode network region. Firstly, the initial kernel was set as the Gaussian function (gray solid line in the figure, mean = 5 sec, standard deviation = 1.5 sec). The shape of extracted HRF throughout the iterations (black dashed line) and the final HRF (color solid line) were illustrated for the synonym task data.

**Figure 5.**
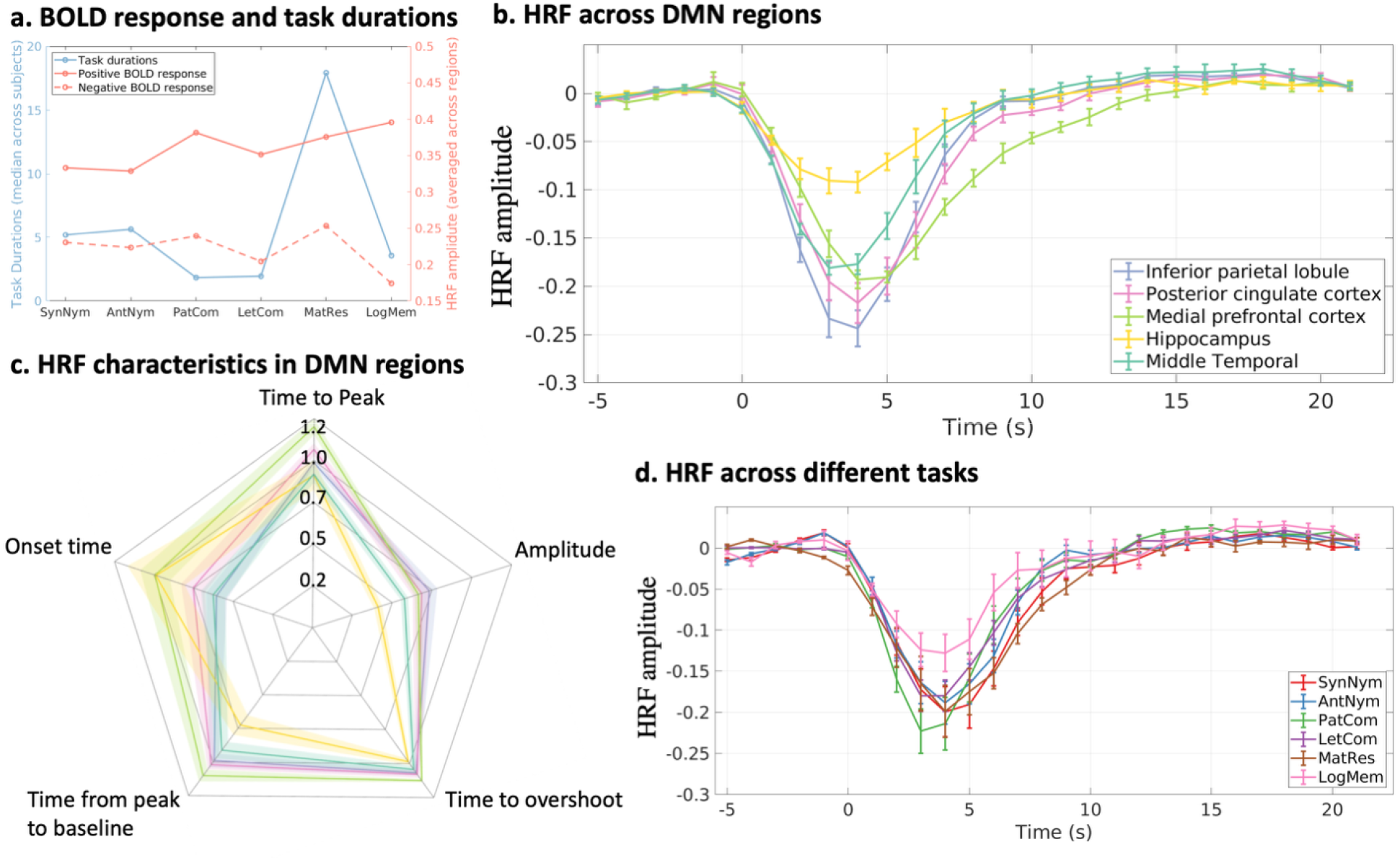
We used iterative finite impulse response (FIR) method to extract hemodynamic response function (HRF) in the default mode network (DMN) regions across six different task fMRI data. (a) Our results demonstrated that the HRFs extracted using our iterative FIR method are not biased by the duration of the stimuli in task design. (b) The averaged HRF across different tasks for each sub-region of the DMN (error bar represents standard deviations across tasks). (c) Various characteristics of the regional HRFs were extracted including onset time, time to peak, amplitude, time to overshoot, and time from peak to baseline, as the ratio to the HRF characteristics of the positive BOLD response. Our results suggest that the HRFs in sub-regions of the DMN exhibit large inter-regional variability and each sub-region has a distinct shape in the response function. (d) The averaged HRF across DMN sub-regions for each task (error bar represents standard deviations across DMN sub-regions). Our results demonstrated a significantly higher inter-regional variability compared to inter-task variability in the HRF of DMN. BOLD, blood-oxygenation-level-depended.

Furthermore, we extracted various characteristics of the regional HRFs including onset time, time to peak, amplitude, time to overshoot, and time from peak to baseline. HRF characteristics in the DMN regions were reported as a ratio over the corresponding characteristic of the HRF of the PBR in the LOC (Figure 5c). Our results suggest that the HRFs in sub-regions of the DMN exhibit large inter-regional variability and each sub-region has a distinct shape in the response function. For example, all the DMN regions have a smaller amplitude compared to the HRF of PBR (IPL: 0.73; PCC: 0.68; MFC: 0.66; Hipp: 0.41; MTG: 0.57). As for the onset time, the HRF of the NBR in the IPL, MTG, and PCC has a faster onset compared to the HRF of the PBR with a ratio of 0.60, 0.62, and 0.75, respectively. And the HRFs of MFC and Hipp have a similar onset time compared to the HRF of the PBR (MFC: 0.99; Hipp: 0.98). The HRFs in the Hipp and MTG reached the peak earlier than the PBR (Hipp: 0.91; MTG: 0.91), whereas the HRFs in the PCC and MFC reached the peak later than the PBR (PCC: 1.06; MFC: 1.20). The HRF in the IPL has a similar time to peak compared to the HRF of PBR (IPL: 0.98). The HRFs in the IPL and PCC have similar time from peak-to baseline to the HRF of PBR (IPL: 0.98; PCC: 1.01). Compared to the HRF of PBR, Hipp and MTG have a HRF with shorter period rising back from the peak to the baseline (Hipp: 0.72; MTG: 0.91), but MFC has a HRF with longer period of returning to baseline (MFC: 1.10). Finally, all the regional HRFs in the DMN have a longer time to undershoot compared to the HRF of PBR (IPL: 1.06; PCC: 1.07; MFC: 1.12; MTG: 1.04), except Hipp demonstrating a similar time to undershoot as the HRF of PBR (Hipp: 0.98).

### Behavioral Analysis

Finally, we assessed the relationship between the HRF amplitude of the NBR in the DMN regions and the behavioral response. For each participant, the behavioral performance was computed as the accuracy over the median response time. In the synonym task, we found that weaker NBR (less negative) in the Hipp is related to better performance (r = 0.300; uncorrected p < 0.027). However, this relationship did not pass FDR multiple comparisons correction, and it was not significant after controlling age and sex (r = 0.2377; uncorrected p < 0.0897). There is no significant relationship between task performance and the HRF amplitude in the other DMN regions (synonym task; IPL: r = −0.056, p > 0.686; PCC: r = 0.086, p > 0.538; MFC: r = −0.049, p > 0.727; MTG: r = 0.036, p > 0.796). Similarly, we did not observe any significant results between NBR HRF amplitude and behavioral response in all other tasks after multiple comparisons correction. Specifically, for the antonym task: IPL: r = −0.025, p > 0.856; PCC: r = 0.178, p > 0.194; MFC: r = −0.066, p > 0.630; Hipp: r = 0.207, p > 0.129; MTG: r = −0.048, p > 0.725. For the pattern comparison task: IPL: r = 0.030, p > 0.825; PCC: r = 0.110, p > 0.417; MFC: r = 0.085, p > 0.528; Hipp: r = −0.079, p > 0.559; MTG: r = 0.033, p > 0.807. For the letter comparison task: IPL: r = −0.030, p > 0.831; PCC: r = −0.022, p > 0.872; MFC: r = −0.138, p > 0.321; Hipp: r = −0.106, p > 0.447; MTG: r = - 0.101, p > 0.468. For the matrix reasoning task: IPL: r = −0.127, p > 0.352; PCC: r = −0.116, p > 0.395; MFC: r = −0.134, p > 0.326; Hipp: r = −0.008, p > 0.952; MTG: r = 0.052, p > 0.706. Finally, for the logical memory task: IPL: r = 0.084, p > 0.555; PCC: r = 0.167, p > 0.238; MFC: r = −0.030, p > 0.831; Hipp: r = 0.132, p > 0.351; MTG: r = −0.045, p > 0.751. Same results were obtained after controlling for age and sex (details in Supplementary Table SI).

## Discussion

Our findings demonstrate that the NBR in DMN regions follows a linear modulation with stimulus duration, and that each region exhibits a distinct HRF shape, which is different from the HRF of the PBR in the visual cortex. Using an iterative FIR approach across multiple cognitive tasks, we found that HRF variability of NBRs is significantly greater between DMN regions than between tasks, with substantial inter-regional differences in amplitude and temporal characteristics. These results provide evidence that DMN subregions have distinct HRF shapes, suggesting region-specific neural and/or vascular mechanisms underlying the NBR in the DMN.

Many studies have provided empirical evidence supporting the existence of the DMN in the human brain, and consistently shown that the medial frontal and medial parietal/posterior cingulate cortex exhibit high resting metabolism, by using fluorodeoxyglucose PET imaging (Raichle et al. 2001). In the resting-state conditions, several fMRI studies using temporal correlations in BOLD signal fluctuations have demonstrated significant within-network functional connectivity of the DMN regions (Greicius et al. 2003; Uddin et al. 2009). Additionally, in task-based fMRI studies with various cognitive tasks, such as recollection of past memories or future-oriented thoughts, robust evoked activations in regions of the DMN were observed (Daselaar et al. 2009; Gusnard et al. 2001; Kjaer, Nowak, and Lou 2002), supporting the involvement of these default mode processes in cognitive tasks.

As the active suppression of activity in the DMN facilitates the execution of cognitive tasks, it is expected that the magnitude of BOLD deactivation increases with task load. Evidence suggests that the intensity of deactivation is modulated by the cognitive load of the task, with greater deactivations in the DMN corresponding to increased task difficulty (McKiernan et al. 2003). For example, in a study using a saccadic task paradigm to investigate error-related deactivations, both error processing and accurate performance were associated with deactivations in the rostral anterior cingulate cortex and other default mode regions (Persson et al. 2007; Polli et al. 2005). Similarly, in another study with a motor-sequence learning task, greater reductions in reaction times during learning were linked to task-related deactivation networks (Tamás Kincses et al. 2008). Furthermore, in a study using a visual global motion detection paradigm, the authors observed a linear increase in task-induced deactivation in the DMN in response to the increase in task difficulty (Singh and Fawcett 2008).

A pioneering study by Lustig and colleagues investigated the NBR in three DMN regions (posterior cingulate, angular gyrus, and medial frontal) during a semantic memory task, where a qualitative comparison of BOLD signals demonstrated the regional heterogeneity between temporal profiles of the NBRs in different DMN regions (Lustig et al. 2003). The MR signal in the posterior cingulate of their study was characterized with a large initial increase before the evoked negative response. In contrast, our results do not show such initial increase in the HRF of the NBR from posterior cingulate. However, time-series analysis obtained using the block-design fMRI paradigms, as in (Lustig et al. 2003), may not provide an accurate representation of the HRF shape (Huettel et al. 2014). To demonstrate the task-dependency of the NBR in DMN regions, Mayer et al. extracted NBR time-series from DMN regions during two distinct tasks involving visual attention and working memory, each with varying cognitive loads. In their study, the extracted time-course in the medial frontal cortex, characterized with a prolonged and extended NBR HRF, is consistent with our findings (Figure 5b). Additionally, in terms of the time-course returning from the negative peak to baseline, the deactivation time-course in the inferior parietal lobule and angular gyrus in our study aligns with Mayer et al., where the HRF raising back to baseline in 6 - 8 seconds. However, the deactivation time-course in the inferior parietal lobule and angular gyrus in our study demonstrated a shorter falling edge from baseline to the negative peak than that reported in Mayer et al. For the posterior cingulate cortex, Mayer et al. reported a sustained deactivation lasting 14 - 22 seconds after the peak before returning to baseline. In contrast, our results indicate a quicker return to baseline from the peak (8 - 9 seconds). Regional dependency in the HRF from the posterior cingulate, medial frontal, and hippocampal sub-regions of the DMN has been previously reported (Meltzer et al. 2008), with an extended delay of 4 - 6 seconds in the onset of the NBR HRF, which is in contrast with our results showing mostly faster or similar onset time in comparison to the HRF of the PBR in the visual cortex.

A more recent study reported a reduction in the magnitude of the NBR during a rule-switching task, where sequential events with identical attentional demands were used (Farooqui and Manly 2018). Deactivations were observed not only in DMN regions but also in other brain regions across these sequential events. Their study also demonstrated different NBR time-courses across various DMN regions. In our study, we employed an iterative FIR approach to detect a robust and reproducible HRF for the NBR across different tasks. Most studies examining the NBR and its spatial and temporal characteristics employ only one or two tasks, where a specific cognitive process was assessed during the fMRI experiment. These tasks are often designed for distinct research objectives and hypotheses. Moreover, our study benefits from advancements in multi-task fMRI acquisitions, providing more reliable results on the extraction of the HRF from the NBR signal in the DMN.

Although emerging evidence has been reported on the neural and vascular mechanisms underlying task-evoked PBR, the mechanisms underlying the accompanying NBR remain largely unknown. As the studies on the NBR have been started since the early days of BOLD-fMRI, little progress has been made on the investigations on the NBR (Allison et al. 2000; Smith, Singh, and Greenlee 2000; Tootell et al. 1998). One potential reason for this lack of progress is that brain regions exhibiting NBR in response to specific tasks may have distinct underlying mechanisms. For instance, the NBR detected in the vicinity (within 10 mm) of the PBR during sensory/motor task stimulation has different properties from the NBR observed in DMN regions, which is typically associated with a decrease in blood flow resulting from suppression of neuronal activity (Ghaderi Yazdi and Razlighi 2025). In line with this, our previous work (He et al. 2022) demonstrated that the HRF of the NBR measured adjacent to PBR regions in the visual cortex differs substantially from the HRFs reported here for DMN regions, both in amplitude and temporal characteristics. This further supports the view that distinct underlying neural and/or vascular mechanisms may generate NBRs in different anatomical regions. Furthermore, the NBR in the ventricles has been linked to changes in cerebral spinal fluid volume (Thomas et al. 2013).

We should emphasize that the present study focuses exclusively on the NBR detected from DMN regions. Therefore, our findings may not be generalized to other types of NBR. A few studies have explored the neural substrates of the NBR in the DMN, demonstrating significant suppression of neural activity in these regions. For example, electrophysiological recordings from the posterior cingulate cortex of macaque monkeys revealed a slight but significant reduction in firing rate during task performance (Pearson et al. 2009). Another study, combining electroencephalography (EEG) and fMRI, found a relationship between NBR in the medial prefrontal cortex and EEG theta rhythm (Mizuhara et al. 2004). A seminal study using stereotactic EEG demonstrated transient neural deactivation, characterized by reduced Gamma-band power in the posterior cingulate and medial prefrontal cortex during task engagement (Jerbi et al. 2010). Together, these findings suggest that neural processes at least partially account for the NBR observed in the DMN, highlighting the importance of understanding the relationship between these processes and the temporal and spatial properties of the NBR in the DMN during task performance.

One of the most challenging aspects of studying the NBR is the technical difficulties and the lack of appropriate methods for investigating this signal. The first challenge arises from the relatively small magnitude of the NBR compared to the PBR, and the second stems from the fact that fMRI processing and analysis methods have been primarily developed for the PBR and are often assumed to be equally applicable to the NBR. For instance, while the linearity of the PBR has been extensively studied (Barch et al. 2003; Birn, Saad, and Bandettini 2001; Boynton et al. 1996b; Dale and Buckner 1997; Huettel and McCarthy 2000; Wager et al. 2005), to our knowledge, the linearity of the NBR with respect to stimulus duration has not been investigated. In this study, we conducted a simple modelling experiment exploring no modulation, linear modulation, and saturated modulation cases for the NBR. Our results suggest that a linear modulation of the NBR provides a better fit to the fMRI data using the synonym task-based fMRI data. As shown in Figure 3, we were able to demonstrate a linear modulation of the NBR with respect to stimulus duration. Most task designs in this study used stimuli lasting longer than one second, and future investigations should focus on shorter stimulus durations, less than one second, which could provide convincing evidence regarding non-linearity or linear modulation in this range. Another technical challenge in studying the NBR is related to the methods used to extract the HRF shape from DMN regions. While numerous deconvolution approaches have been proposed for the PBR (Glover 1999; Goutte et al. 2000; Josephs, Turner, and Friston 1997; Woolrich, Behrens, and Smith 2004), their applicability to the NBR has not been adequately evaluated. To avoid imposing prior assumptions on the NBR HRF shape, we used the FIR method for HRF extraction (Goutte et al. 2000). Initially, we selected voxels from DMN regions that showed significant deactivation with a starting Gaussian function kernel. However, by using this preliminary voxel selection approach, the extracted HRF will be biased towards the kernel in the GLM analysis. To avoid this, we developed an iterative FIR method, which refines the voxel selection based on the extracted HRF from the previous iteration, iterating until convergence to the final HRF shape. We validated the iFIR method on multiple datasets with various cognitive tasks and demonstrated that even starting with a Gaussian kernel, the iFIR converges to a similar shape HRF for each region in the DMN. When examining associations between HRF amplitude and behavior response, we found no significant effects across DMN subregions or across tasks. This result aligns with our expectation that the HRF, as a unit response, should be invariant to behavioral performance. Our results suggest that the NBR HRFs estimated with our iFIR method do not appear to be biased by participants’ performance.

A growing body of literature reports alterations in NBR within the DMN across both healthy and clinical populations (Ghaderi Yazdi and Razlighi 2025; Jacobs et al. 2014; Persson et al. 2007). For instance, clinical conditions such as Alzheimer’s disease (AD), schizophrenia, autism, attention deficit hyperactivity disorder (ADHD), and other mental disorders (Assaf et al. 2010; Broyd et al. 2009; Buckner, Andrews-Hanna, and Schacter 2008; Greicius et al. 2004; Kennedy, Redcay, and Courchesne 2006; Liu et al. 2006; Lustig et al. 2003; Petrella et al. 2007; Rombouts et al. 2005) have been associated with NBR changes in DMN regions. Furthermore, differential NBR can be mistaken as PBR in second-level analyses, as previously noted (Gusnard et al. 2001), potentially contributing to an underreporting of NBR findings. Interventional studies have also highlighted the role of NBR. For example, in a medication study with healthy adults, an increase in NBR magnitude was observed in the posterior cingulate and medial prefrontal cortex in this group compared to controls (Seibert and Brewer 2011). Lustig and colleagues demonstrated that NBR in the posterior cingulate could differentiate demented patients not only from healthy young adults but also from age-matched healthy older adults (Lustig et al. 2003). In patients with AD, a global reduction in NBR magnitude was reported across all DMN regions (Persson et al. 2007), while in subjects with age-related memory impairment, the reduction was restricted to the parietal area, with no alteration observed in the hippocampus (Miller et al. 2008). Furthermore, studies of participants with amyloid deposition and APOE4 carrier status also showed reduced NBR magnitude in DMN regions, similar to patterns seen in AD patients, suggesting that NBR could serve as an early biomarker for AD in asymptomatic individuals at higher risk (Hafkemeijer, van der Grond, and Rombouts 2012). Alterations in NBR have also been reported in patients with depression, particularly in the hippocampus, parahippocampus, and amygdala, which may contribute to the inability to control negative thoughts and emotions (Sheline et al. 2009). Neuropsychiatric disorders with cognitive impairment, such as major depressive disorder and schizophrenia, have also been shown to affect NBR in the DMN (Pomarol-Clotet et al. 2008; Sheline et al. 2009). For example, alterations in NBR in the prefrontal cortex, including the anterior cingulate and gyrus rectus, are thought to underlie impaired cognitive functions and symptoms such as disorganized speech, inability to concentrate, and difficulty maintaining a train of thought in schizophrenia patients (Andrews-Hanna, Smallwood, and Spreng 2014; Pomarol-Clotet et al. 2008). Lastly, an intriguing study on disorders of consciousness demonstrated that NBR in the DMN can differentiate between patients with minimal consciousness syndrome and those with unresponsive wakefulness syndrome, the latter showing no detectable NBR in the DMN (Crone et al. 2011). Our findings introduce a potential confounding factor for studies using the same HRF for NBR as for PBR, which could lead to false-positive results regarding regional heterogeneity. Therefore, we recommend that future studies investigating NBR should adopt region-specific HRF kernels for DMN regions, as identified in this study, to avoid such confounding effects.

Each region of the DMN encompasses a large portion of the brain, often extending into multiple neuroanatomical areas with distinct functions, neuronal cytology and densities. Therefore, it is plausible to hypothesize that each region may have more than one functionality, each associated with a unique HRF tightly coupled to its underlying neural processes. Preliminary evidence of within-region functional and structural heterogeneity in the DMN has been reported for the posterior cingulate and lateral parietal regions (Leech et al. 2011; Shulman et al. 2003, 2007; Vogt, Vogt, and Laureys 2006). This suggests the possibility that each DMN region could potentially exhibit multiple HRF kernels, reflecting parallel processes that contribute together to the final shape of the NBR HRF. We explored this possibility in our dataset using voxel-wise HRF clustering analysis within each region. In a few subjects, we observed more than one HRF shape in the posterior cingulate. Voxels near the brain surface and large arteries exhibited a fast, biphasic kernel with relatively larger magnitude, consistent with previous findings (Meltzer et al. 2008). In contrast, voxels deeper in the ventral part of the region showed a slower kernel with greater emphasis on the negative phase, more closely to the shape of the final HRF results reported in this study. While this observation was intriguing, we were unable to replicate it in most of our subjects, leading us to exclude it from the main results. However, future studies with higher signal-to-noise ratios, focusing primarily on the posterior cingulate, may help to further disentangle this observation.

While most of the existing studies show that NBR in the DMN are equivalently observed in all sort of cognitive and sensory-motor tasks independent of the type of the task being performed, there are few studies highlighting the possibility of the task-specificity of the NBR in the DMN regions (Mayer et al. 2010; Shulman et al. 2007). Furthermore, we have also shown previously using regional activation maps (Parker and Razlighi 2019) and in this work using the extracted HRF that the NBR in the DMN is attention-specific. Therefore, it is also possible that different tasks may give rise to different HRF kernel in each region of the DMN, which in turn itself can generate regional heterogeneity in the NBR obtained from different regions, if the differences in the HRF shapes are not taken into account. In this study, we used various tasks to extract the HRF with the aim of extracting the most general HRF shape possible for the NBR. Our results demonstrated a significantly greater inter-regional variability than inter-task variability in the HRF of DMN. However future investigation is warranted to study whether different types of tasks (e.g. working memory versus perceptual speed) will significantly alter the shape of the HRF in the DMN regions.

## Conclusion

As the utilization of the NBR in investigating brain-behaviour relationships as well as neurological and psychiatric conditions becomes more popular, the need for complete understanding and characterization of the NBR becomes essential to be able to accurately interpret the results of such studies. In this study, our results provided evidence supporting the linearity assumption of task modulation in the DMN regions. We developed and used an iterative FIR approach to extract the NBR HRF kernel in each region of the DMN, demonstrating that each region presents distinct HRF kernel, which is distinct from the HRF of the PBR in the visual cortex. The results highlight the possibility of having unique underlying neural and/or vascular mechanism specific to the NBR in each DMN region.

